# Watching Yourself Talk: Motor Experience Sharpens Sensitivity to Gesture-Speech Asynchrony

**DOI:** 10.64898/2026.02.12.705486

**Authors:** Tiziana Vercillo, Judith Holler, Uta Noppeney

## Abstract

Language is inherently multisensory, with speech often accompanied by iconic gestures that convey semantic meaning related to actions, objects, or spatial relationships. Although the temporal coordination between speech and gesture is variable, the brain integrates these signals seamlessly. Yet, much of the cognitive mechanisms behind this integration remain unclear. This study investigates whether sensorimotor experience, specifically with one’s own speech and gestures, enhances temporal sensitivity through internal forward models that guide audiovisual prediction. Participants first produced sentences with corresponding iconic gestures, which were audiovisually recorded. These recordings were later temporally manipulated and presented in a simultaneity judgment task, where participants evaluated both their own and others’ recordings. Results revealed narrower temporal binding windows (TBWs), indicating heightened sensitivity to audiovisual asynchrony, when participants judged their own speech-gesture recordings compared to those of others. To further explore the role of motor experience, we analysed individual variability in gesture-speech timing during production and found no reliable relationship between production variability and perceptual sensitivity, suggesting that perceptual precision is not simply a reflection of motor consistency. These findings demonstrate that sensorimotor experience with self-generated movements sharpens multisensory temporal integration, likely via predictive internal models, and underscore the functional role of predictive motor mechanisms in supporting temporal integration across perceptual and action systems.

## 1. Introduction

Communication is a fundamental human behaviour, yet many underlying cognitive mechanisms remain elusive. Traditionally, language has been considered a primarily verbal activity, but in natural face-to-face interactions, it is inherently multimodal (Holler & Levinson, 2019; McNeill, 1992; Vigliocco et al., 2014). Spoken words are frequently accompanied by hand gestures, which carry semantic and pragmatic meaning. Iconic gestures, in particular, visually depict actions, objects, or spatial relationships, and are integrated with speech during comprehension (Dargue et al., 2019; Drijvers & Özyürek, 2017; Holle et al., 2010; Kelly et al., 2004; Kelly, Özyürek, et al., 2010; Momsen et al., 2021). For example, when a speaker talks about cooking while simultaneously making a flipping motion with their hand, the gesture and the spoken word “cooking” are bound together, enhancing the listener’s understanding. This is particularly useful since iconic gestures often add visual elements that complement speech, enriching the overall cognitive representation. In this case, the flipping gesture provides additional semantic information, allowing the listener to infer that the speaker is specifically referring to pan-frying rather than another cooking method.

Unlike other multisensory events (Holmes & Spence, 2005; Stein et al., 2014; Stevenson et al., 2012; Mark T. Wallace & Stevenson, 2014), the temporal alignment of speech and gesture is highly variable, such that the two events rarely unfold in precise synchrony. Gestures often precede their corresponding words, with variability influenced by communicative context and speaker style (De Jonge-Hoekstra et al., 2021; Obermeier & Gunter, 2014). On average, gestures seem to begin ∼644 ms before their lexical affiliates (the lexical items they most closely relate to semantically), with the most meaningful gesture phase (the stroke) occurring ∼193 ms before the spoken word (ter Bekke, Drijvers & Holler 2024). Nevertheless, despite this considerable asynchrony, the brain readily binds speech and gesture. This raises the question of which factors support the integration of speech and gesture despite their substantial temporal variability.

One way to address this question is to consider findings from the broader multisensory integration literature. Evidence from this work suggests that temporal sensitivity is fundamental for the integration of auditory and visual stimuli. For instance, a narrower temporal binding windows (TBW) is associated with a better performance in multimodal speech perception and to a better understanding of speech in noise (Kunnath et al., 2025; Zerr et al., 2019). Moreover, across a range of neurodevelopmental disorders, including autism, dyslexia, and schizophrenia, an enlarged TBW has been linked to deficits in multisensory integration and broader cognitive functioning (Wallace & Stevenson, 2014). Importantly, the relationship between temporal processing and multisensory integration generalizes across stimulus domains, extending from simple artificial stimuli to more complex, naturalistic contexts such as music and language. Together, these findings suggest that temporal sensitivity may be crucial for the integration of gesture and speech, and raise the possibility that factors shaping temporal sensitivity play a key role in audiovisual language processing.

Consistent with this view, a previous study by Lee & Noppeney (2011) suggested that sensorimotor experience is a critical factor in shaping audiovisual temporal binding. In their study, expert musicians demonstrated greater sensitivity to audiovisual asynchronies in music-related stimuli compared to non-musicians. Precisely, musicians showed a narrower TBW for audiovisual music stimuli, i.e. a smaller temporal interval within which they perceived the two events as occurring together, making them more sensitive to the smallest temporal delay. These behavioural differences were mirrored by changes in neural activity within the superior temporal sulcus–premotor–cerebellar circuitry, linking audiovisual temporal integration to action processing. However, the study used a between-subjects design focused on piano practice, leaving open questions about potential confounds such as individual differences in temporal sensitivity and did not isolate the role of familiarity with specific sensorimotor associations from broader expertise effects. Furthermore, music and speech differ fundamentally in their cognitive and neural processing demands. While both share temporal structures, only speech relies on semantics.

We propose that individuals’ familiarity with their own movements and speech increases their sensitivity to subtle temporal discrepancies between co-occurring gesture and speech signals, leading to a narrower temporal binding window (TBW). This idea aligns with theoretical frameworks proposing that internal forward models, shaped by one’s experience, enhance temporal prediction and thereby improve the perception of corresponding multimodal signals (Fincher-Kiefer, 2019; Friston, 2010; D M Wolpert et al., 1995). We argue that the characteristic patterns of one’s own audiovisual speech refine these predictive mechanisms and, in turn, heighten sensitivity to the (a)synchrony between gesture and speech, which may affect semantic integration.

The TBW, which defines the temporal limits for multisensory integration, is known to be influenced by stimulus complexity, individual differences, and experience-dependent plasticity (Stevenson & Wallace, 2013; Van Eijk et al., 2008; Vatakis & Spence, 2006; Wallace & Stevenson, 2014b). It is broader for naturalistic stimuli like lip movements and speech than for simple flashes and beeps and can be shaped through recalibration or training. Here, we suggest that sensorimotor experience not only modulates the TBW but does so in a self-specific manner. We hypothesize that individuals develop heightened sensitivity to temporal misalignments in their own gesture-speech patterns compared to those of others. This would suggest that multimodal integration is not solely driven by general perceptual principles but is also shaped by individualized sensorimotor experiences, emphasizing the dynamic interplay between action, perception, and prediction in communication.

## 2. Methods and procedures

### 2.1. Participants and recording protocol

The study initially included 24 individuals; however, two of them did not complete all the required sessions, and one participant was not considered in the behavioural analysis because they consistently selected the same response in the two-alternative forced-choice (2AFC) task (see below), suggesting an inability to engage meaningfully with the task requirements. Overall, we had a total of 21 participants included in the final analysis. All the participants were Dutch native speakers and had English as a second language, had normal or corrected-to-normal visual acuity and normal hearing, and did not report any speech or motor impairments. All the participants gave written, informed consent prior to participating in the study. The experimental protocol was approved by the Social Science Faculty Ethics Committee at Radboud University, Nijmegen, The Netherlands.

The study included two separate experimental sessions: “production” and “perception”. The production session was consistently scheduled first, as its primary objective was to “produce” audio-visual recordings of the participants, which would subsequently serve as stimuli for the perception experiment, involving the same participants as in the production sessions, except for one participant who did not participate in the perception task.

### 2.2. Experimental Design: From Production to Perception

#### 2.2.1. Production

This experimental session was held in a soundproof room where participants took part in the production session individually. Participants were standing while a video camera recorded their body from a frontal position (Canon XF205 Camcorder) and high-quality audio was recorded using a Sennheiser ME64 microphone. Audio and video were synchronized using Metus INGEST and Adobe Premiere Pro CS6. Videos were recorded at a rate of 25 frames per second, providing a temporal resolution of 40 milliseconds per frame.

A written list of 10 Dutch sentence stems was created, each with two possible endings containing an action verb, resulting in a total of 20 different sentences. These stems typically occurred at the end of a more complex structure that included an initial contextualizing clause. For example, one sentence might begin as follows: “*Ik kook Pastasaus. Eerst moet ik de tomaten…*” with possible completions being either “*bakken*” or “*snijden*” meaning “*I’m cooking pasta sauce. First, I need to <u>fry/cut</u> the tomatoes*”. Participants were first given time to familiarize themselves with the list of sentences. They were then instructed to say each sentence aloud in a natural manner and to accompany the final word with a hand gesture that represented its meaning (thus rendering the final word the ‘lexical affiliate’ of the gesture, defined as the lexical item in the speech that is most closely related to the semantic depiction of the gesture). Importantly, these gestures were not scripted but were produced spontaneously by the participants. To examine within-subject variability in the temporal production of speech and gesture, participants completed 10 repetitions of each sentence. This produced 200 audiovisual recordings (20 sentences × 10 repetitions), collected across 10 blocks of roughly 15 minutes each, with each block containing one full set of the 20 sentences.

#### 1.2.1. Gesture - Speech production: Stimulus Manipulation and Asynchrony Conditions

Four sentences were selected from the original list to serve as stimuli for the perceptual experiment (see Supplemental Material). Selection was based on cross-participant consistency in gesture production, specifically the use of the same hand and comparable movement trajectories, which resulted in gestures with closely aligned kinematic profiles. From the ten repetitions participants produced for each sentence, one video per sentence was chosen for each participant. This resulted in four videos per participant and a total of 96 videos across all participants.

Videos were edited using the software DaVinci Resolve 20. To ensure partial anonymity, participants’ faces were blurred, while their voices remained unaltered. All participants consented to the use of their recordings as stimuli, despite the absence of complete anonymity due to the retention of bodily information and voice identity.

Eleven stimulus onset asynchronies (SOAs) were created for each video. Audio-visual asynchronies were produced by shifting the onset of the whole audio track relative to the onset of the whole video track, plus inserting static video and audio frames (ambient noise) at either the beginning or the end of each original video to compensate for the artificially introduced delays (see Figure 1 for a graphical representation of the procedure). The SOAs used in the study were: ±800; ±640; ±480; ±320; ±160; 0 ms, with negative values indicating that the action verb preceded the gesture (audio leads) and positive values indicating that the gesture preceded the action verb (vision leads). See Figure 1A for a graphic description of the stimuli manipulation.

**Figure 1.**
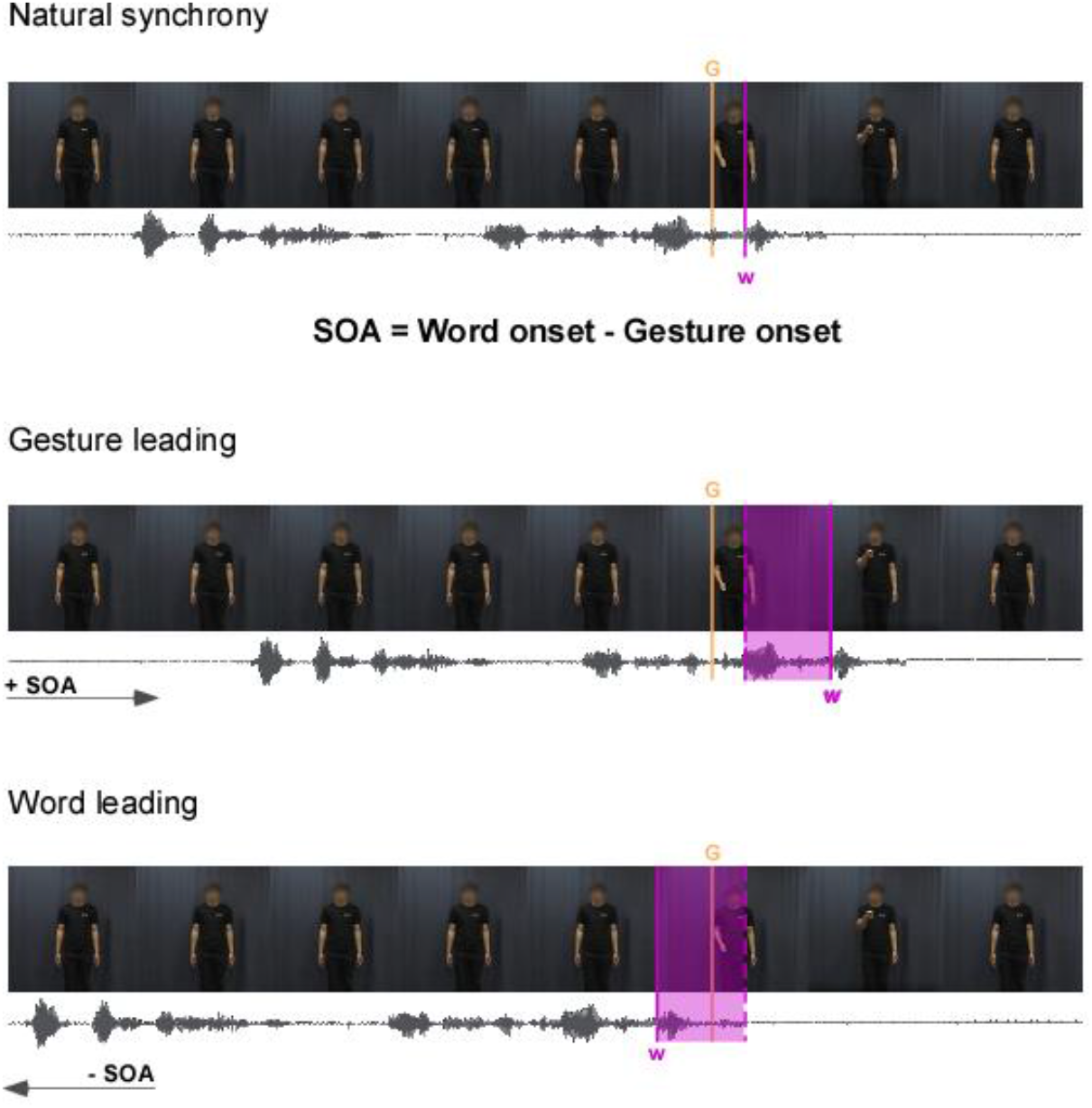
Graphic illustration of the audio-video manipulation used to generate asynchrony values. The top panel shows the original video and the natural temporal relationship between the onset of the gesture (G) and the word (W). The middle panel illustrates a positive SOA condition, where the audio track is shifted backward, delaying the onset of the word and increasing the asynchrony between the word and the gesture with the gesture preceding the target word (i.e. visual leading). The bottom panel depicts the opposite condition, with a negative SOA value, where the audio track is shifted forward, causing the target word to occur earlier and reversing the natural synchrony between the word and the gesture (audio leading).

#### 1.2.2. Gesture – Speech Perception

This session constituted the main experiment. For analytical purposes, participants were grouped into 12 pseudo-pairs so that each participant was presented with stimuli generated from both their own recordings (referred to as the SELF condition) and those of the respective other participant within the pair (the OTHER condition). Participants in each pair were always strangers to one another. To assess participants’ perception of audiovisual synchrony in these speech stimuli, the Simultaneity Judgment (SJ) task was employed.

In the SJ task a series of videos with different SOA values were presented on a computer screen and the participant was required to indicate, upon video completion, whether the gesture and the final word were perceived as synchronous or not by pressing a designated key on the keyboard (refer to Figure 1B for a visual representation of the task). Participants were specifically instructed that there was only one gesture per sentence and that the gesture always referred to the final word of the sentence. The SJ task is commonly used in psychophysical experiments to measure the TBW for audiovisual stimuli (Stone et al., 2001; Zampini et al., 2005).

Before starting the experiment, participants underwent a familiarization phase, where they were introduced to the concept of “natural” synchrony. This concept illustrates how, within a movie, gestures and words occur together, in a cohesive manner. We further showed them how the same movie would look like after temporal manipulation at the most extreme SOA values. The demonstration used recorded movies that did not feature any of the participants within the participant pairs. The experiment was implemented using Python and PsychoPy. Participants completed 13 blocks in total. Each block consisted of trials from the two conditions: SELF and OTHER, presented separately. The order of these conditions was randomized across blocks and participants. Within each block, each condition included 44 trials, consisting of 4 sentences combined with 11 different SOA values, presented in a random order. At the end of each video, a question appeared on the computer screen prompting participants to make a simultaneity judgment by responding through button press. Overall, participants completed a total of 572 trials per condition (4 sentences x 11 SOAs x 13 blocks) and given the two conditions (SELF/OTHER), 1144 trials in total.

### 1.3. Data Analysis

#### 2.3.1 Measuring Perceptual Sensitivity and Temporal Binding

Individual data were collapsed across blocks but analysed separately for the different sentences and experimental conditions (SELF vs OTHER). For each of the 4 sentences x 2 conditions, we measured the proportion of trials where the word and the gesture were perceived as simultaneous at each SOA value. Such individual proportions were then fitted with Gaussian distributions using the MATLAB function ‘fit’ (with fittype = ‘gauss1’) to estimate: 1) the stimulus- and participant-specific Point of Subjective Simultaneity (PSS), i.e. the mean of the Gaussian distribution - representing the asynchrony value at which the participant perceived the word and the gesture as being synchronous most of the time; 2) the Temporal Binding Window (TBW), i.e. the standard deviation of the Gaussian distribution, representing the overall time interval within which the two sensory events are likely perceived as synchronous.

Because the temporal binding window (TBW) is derived from perceptual judgments, it may be influenced not only by temporal sensitivity but also by direction-dependent biases or shifts in decision criterion. Assuming a symmetric TBW imposes the constraint that such biases are equivalent for estimates obtained when the visual stimulus leads versus when the auditory stimulus leads the audiovisual event. It is therefore important to explicitly test for asymmetry in the shape of the TBW. To assess whether allowing asymmetry improved model fit, we compared a symmetric Gaussian model to an asymmetric split-Gaussian model. Both models were fit to each dataset using nonlinear least-squares optimization. Model comparison was performed using the Bayesian Information Criterion (BIC), which balances goodness of fit against model complexity. Differences in BIC (ΔBIC = BIC_sym_ - BIC_asym_) were used to evaluate evidence for the asymmetric model, with ΔBIC values smaller than 2 taken as negligible evidence for improvement and values greater than 6 indicating strong evidence. As a confirmatory analysis, likelihood-ratio tests were also conducted to compare the two models. Allowing asymmetric widths did not result in a meaningful improvement in model fit. Across subjects and conditions, ΔBIC values clustered near zero (ΔBIC_self_ = -1.49, ΔBIC_other_ = -1.20), and likelihood-ratio tests showed no significant advantage for the asymmetric model (p = 0.46).

The first hypothesis tested whether participants’ temporal binding window (TBW) was narrower in the SELF condition compared to the OTHER condition. In other words, we examined whether participants were more sensitive to audiovisual asynchronies in their own recordings than in those of a stranger. To this aim, TBW values were compared using a linear mixed-effects model with maximum likelihood estimation (MLE). The full model was specified as:

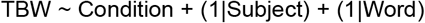

with *Condition* (SELF vs. OTHER) included as a fixed effect, and random intercepts included for *Subject* and *Word*. To assess the contribution of condition, the full model was compared to a reduced model excluding the fixed effect:

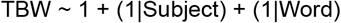

After selecting the best-fitting fixed-effects structure, the final model parameters were estimated using restricted maximum likelihood (REML).

The same modelling approach was applied to PSS, to assess whether participants’ PSS varied across conditions. Model comparisons were again conducted using likelihood ratio tests on models fitted with maximum likelihood estimation.

#### 2.3.2 Linking variability in production to perceptual sensitivity

After examining how sensorimotor experience (reflected in the SELF/OTHER conditions) influenced the TBW, we investigated whether individual differences in speech motor patterns contributed to this effect. Motor production plays a central role in shaping sensorimotor experience, as the timing and coordination of articulatory movements provide internal feedback that supports the prediction and integration of sensory input. We therefore aimed to determine whether the temporal dynamics of speech production varied systematically across individuals, potentially accounting for individual differences in audiovisual perception. To address this question, we measured individual variability in the timing of gesture–speech production and examined whether it was related to individual differences in perceptual TBW, separately for each condition and word.

Videos selected in the production session were analysed using a frame-by-frame method to identify the onset of the gesture as a whole movement. The gesture onset was defined as the first frame in which the hand(s) began moving away from their rest position and appeared blurred (ter Bekke et al., 2024). For each gesture, the onset of the target word was determined from the audio. This was the first noticeable spike in the waveform representing the silent-to-sound transition, measured through the software Elan 5.9 and Praat (version 6.1.15). The difference between word onset time and gesture onset time for each recording was measured to assess the degree of “natural” synchrony between speech and gestures. Individual variability in sensorimotor production was computed for each gesture-word pairing as the standard deviation of gesture-word natural synchronies across repetitions of each sentence.

To examine whether the variability in the timing of gesture-speech production predicts perceptual variability (i.e. the TBW), we used linear mixed-effects models including production variability (ProdVar) and condition (self/other) as fixed factor, and their interaction (ProdVar × Condition). Random intercepts were included to account for variability across participants and words. Once again we compared a full model (fixed effects + random effects):

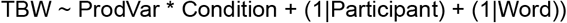

to a reduced model without the effect of production variability:

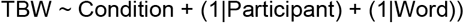

Model comparisons were conducted using likelihood ratio tests on models fitted with maximum likelihood (ML). The final model, including significant effects, was refitted using restricted maximum likelihood (REML) to obtain unbiased estimates.

## 2. Results

The three hypotheses we tested were: (1) that participants would be more sensitive to temporal asynchrony in the SELF condition than in the OTHER condition, reflected in a narrower TBW; (2) that accuracy would differ between conditions, reflected in a shift in the PSS; and (3) that variability in gesture–speech timing during production would predict variability in gesture–speech timing perception.

Figure 2 presents the average performance in the SJ task across the two experimental conditions (SELF vs. OTHER) for each of the four words. For each SOA value, we calculated the proportion of ‘simultaneous’ responses per participant and then averaged these across participants. Although the primary purpose of this figure is to visualize the model fitting and overall data distribution, it is already apparent that the OTHER (grey) distribution exhibits a greater standard deviation, indicating a larger TBW, than the SELF (black) distribution, and that the two distributions overall have similar means, with little shift in the PSS.

**Figure 2.**
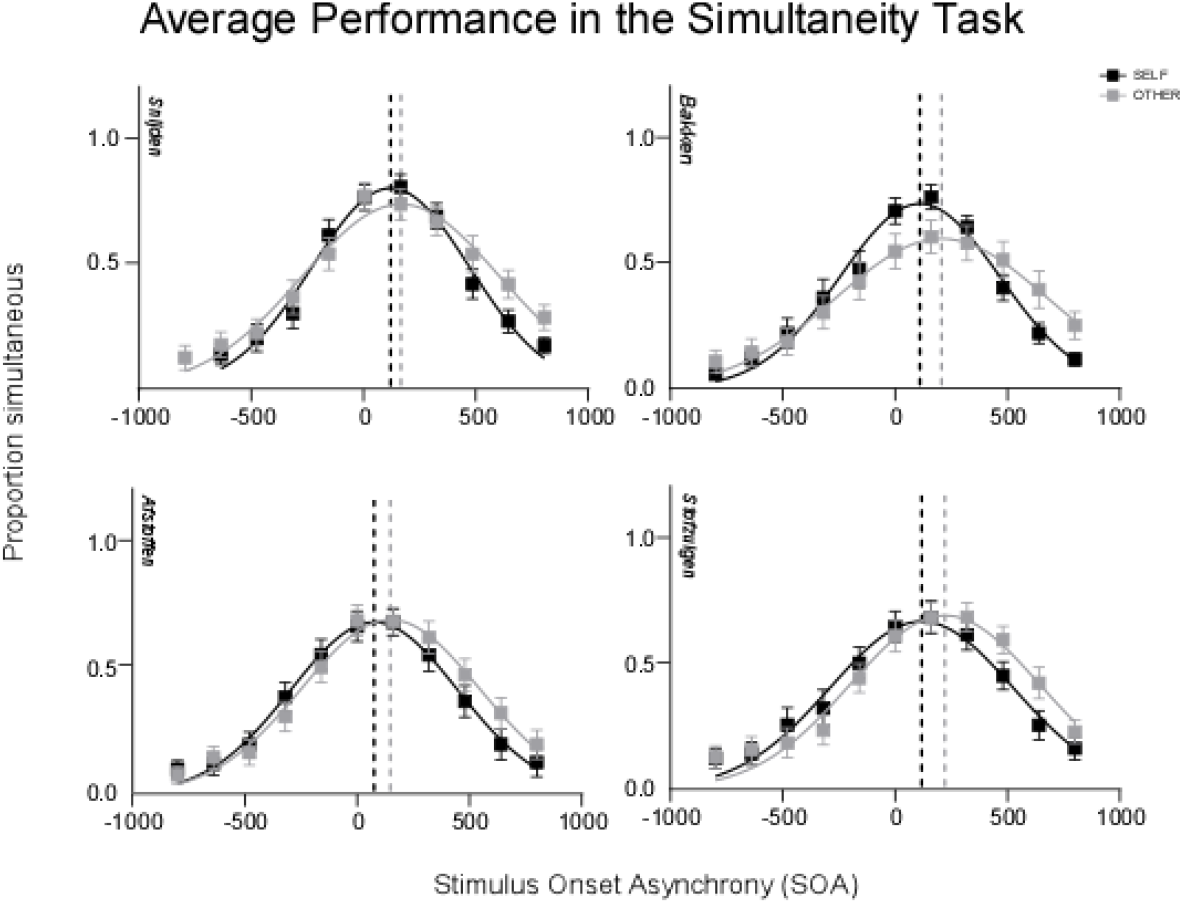
Group averages for the simultaneity tasks are shown for each of the four words in the SELF (black) and OTHER (grey) conditions. Symbols represent the group mean ± standard error at each SOA value, and the lines represent Gaussian fits to the data. Dashed lines mark the means of the fitted distributions. An SOA of 0 indicates perfect simultaneity between the gesture and the word; positive SOA values indicate the gesture preceded the word, while negative SOA values indicate the word preceded the gesture.

For a more detailed analysis, individual data were fitted with a Gaussian function to extract the two key parameters: the Temporal Binding Window (TBW), represented by the curve’s standard deviation, and the Point of Subjective Simultaneity (PSS), corresponding to the curve’s mean. These individual TBW and PSS values were then compared across conditions, as detailed below.

### 3.1 Self-Generated Speech-Gesture Utterances Lead to a Narrower Temporal Binding Window

After comparing the full model (fixed effect + random effects) to a reduced model (random effects only) using MLE we found that the fixed effect of condition (SELF vs. OTHER) significantly improved model fit (χ^2^(1) = 10.35, p = .0013). The final model was therefore refitted using REML. As hypothesized, the results revealed a significant effect of condition on TBW (β = 49.82, SE = 15.21, t(166) = 3.27, p = 0.001), indicating that participants were more sensitive to asynchronies when evaluating their own speech-gesture recordings as opposed to those of another person. In Figure 3, we display TBW values across participants, target words, and conditions.

**Figure 3.**
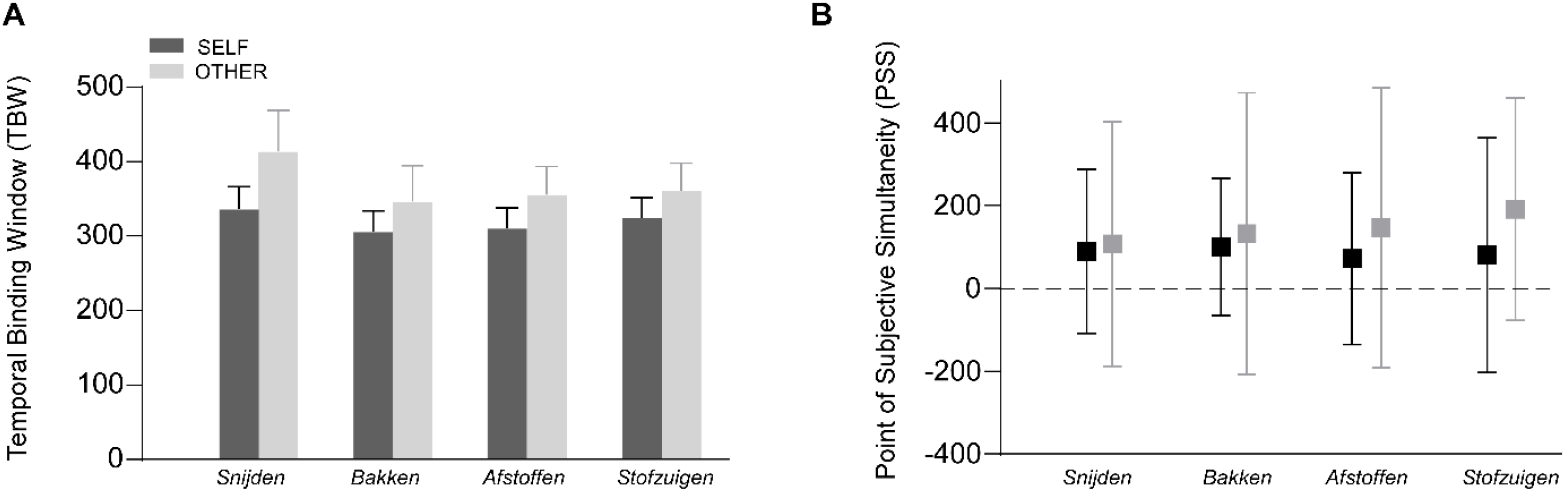
A) Individual variations in the TBW (left panel) B) and PSS (right panel) across the different target words (snijden, bakken, afstoffen, and stofzuigen) for the SELF (black) and OTHER (grey) conditions. Each floating bar starts at a lower bound (i.e. minimum value) and ends at an upper bound (i.e. max value), visually representing the difference of range between conditions. The lines within each bar represent the group average, providing a visual reference for central tendency.

### 3.2 No Shift in Perceived Subjective Simultaneity

In our second hypothesis we examined whether participants’ perceived simultaneity differed when observing self-generated and other-generated speech-gesture utterances. A likelihood ratio test comparing the full model (fixed effect of condition + random effects) to a reduced model (random effects only) did not indicate a significant improvement in model fit (χ^2^(1) = 3.22, p = .073). This result suggests that participants’ sensitivity to simultaneity was not systematically influenced by whether they were evaluating their own recordings or those of another person. However, the model indicated that intra-participant variation played a significant role in perceived simultaneity, highlighting individual differences in temporal processing. In Figure 3 (right panel), we illustrate individual variability and group average PSS across participants, target words, and condition.

Interestingly, all participants exhibited a positive bias, likely reflecting the natural tendency for gestures to precede their lexical affiliates in time. To further investigate this, we explored the relationship between production and perception. Specifically, we tested whether variability in gesture-speech timing during production could account for uncertainty in gesture-speech timing perception.

### 3.3. Production Variability Predict Perceptual Precision only in the other condition

Figure 4A displays individual natural synchrony values, separated by word and averaged across the 10 blocks. While all values are positive, indicating that gestures consistently precede their lexical affiliates, there is notable individual variability in speech-gesture timing. This variability is further illustrated in Figure 4B, which shows the standard deviation of gesture-speech timing across the 10 blocks for each participant.

**Figure 4.**
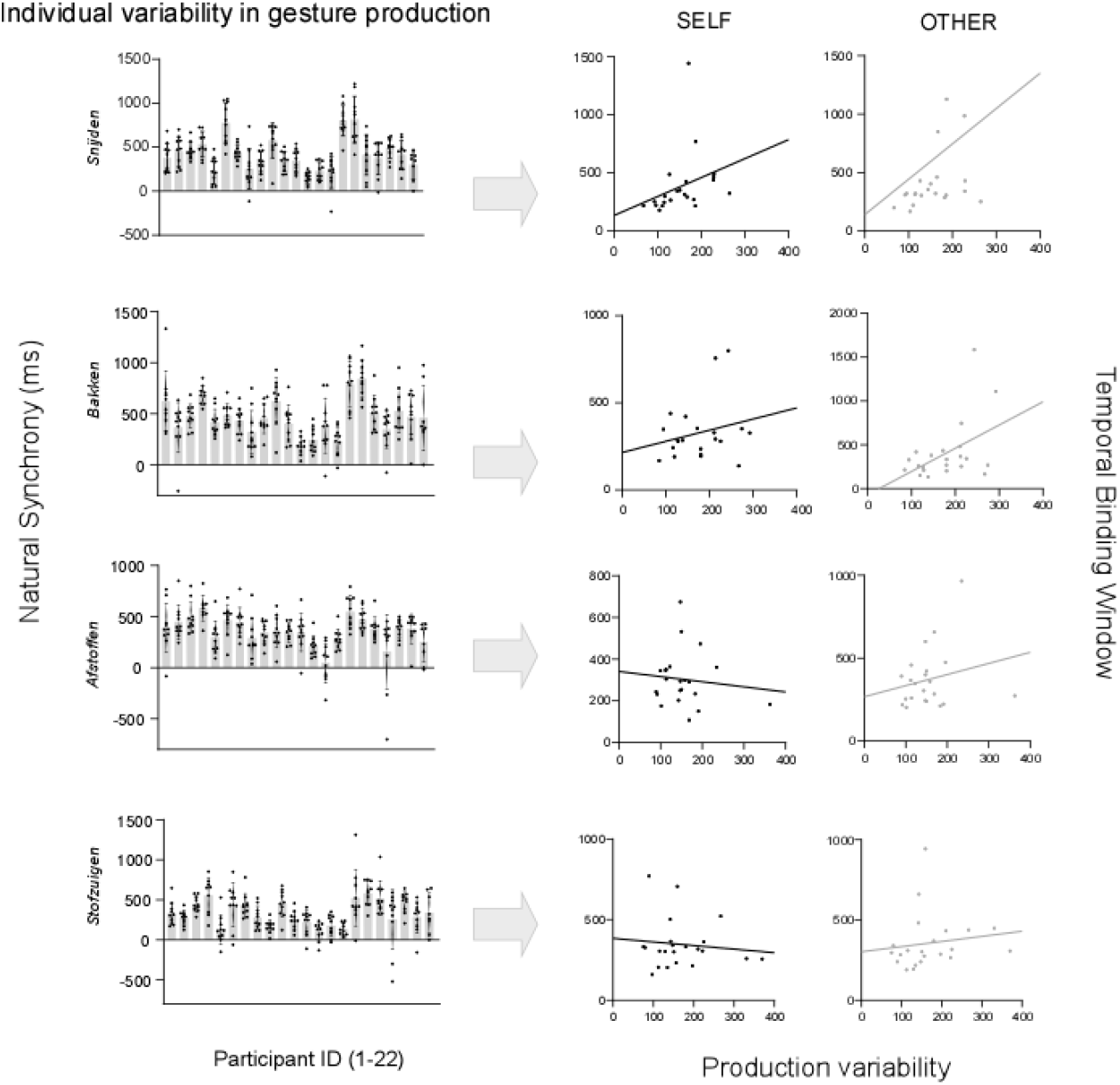
The left panels show the natural synchrony between the gesture and the word for each of the four target words, displayed separately for each of the 22 participants. Each symbol represents the participant’s average (± standard error) across 10 blocks. Individual trials are represented by the black dots. The right panels illustrate the correlation between individual variability in gesture-speech timing (x-axis) and the individual TBW (y-axis), for the SELF and OTHER conditions respectively.

Including production variability as a fixed effect significantly improved model fit relative to the reduced model (χ^2^(2) = 8.89, p = .012). In the final model fitted with REML, the interaction between condition and production variability was significant (β = 0.63, SE = 0.24, t(164) = 2.67, p = .0084), indicating that the relationship between motor and perceptual variability differed across conditions. Specifically, higher production variability was associated with larger TBW when participants observed another person, whereas this relationship was weaker when observing themselves. These results suggest that motor variability influences perceptual variability in a condition-dependent manner.

To explore the link between individual variability in the production and perception of gesture-speech timing, we computed correlations between participants’ variability in natural gesture timing and their temporal sensitivity in the SJ task (i.e., TBW), separately for each word and condition. In the SELF condition, no significant correlations were found for any of the words. In the OTHER condition, a significant positive correlation emerged only for the word “bakken” (r=0.47; p=0.027). These results suggest a potential relationship between individual differences in gesture-speech timing during production and perceptual sensitivity, but only under the OTHER condition and for one of the four lexical items used as stimuli.

## 3. Discussion

The present study aimed to investigate how sensorimotor experience influences the perception of temporal alignment between speech and iconic hand gestures. Our findings revealed two key insights. First, the temporal integration of spoken words and hand gestures was modulated by sensorimotor experience. This was supported by the finding that familiarity with one’s own speech–gesture utterances improved the ability to detect temporal mismatches between words and gestures, likely through the development of robust internal predictions. Second, variability in the timing of gesture production during speech influenced temporal sensitivity in gesture-speech binding, but this effect emerged only when observers viewed an unfamiliar speaker and was limited to a specific lexical affiliate. In other words, individuals who produced gestures with highly consistent timing show greater sensitivity to temporal offsets between speech and gesture when observing others speak. Together these results support the critical role of internal forward models, formed through repeated experience with self-generated actions, in shaping multimodal perception and support the idea that timing mechanisms are shared between motor and perceptual systems.

In the context of multimodal communication, internal models (Friston, 2010; Wolpert et al., 1995), may play a crucial role in predicting the temporal occurrence of words and gestures and estimate their temporal alignment resulting in more effective integration. Here, we argue that enhanced temporal processing arises from the predictive capabilities of internal models and is shaped by familiarity with speech-gesture utterances. The sensorimotor experience inherent in daily communication likely refines these internal models, strengthening the brain’s ability to anticipate the timing of familiar speech-gesture combinations and contributing to the greater sensitivity to temporal asynchronies observed in our study. And even for unfamiliar gesture-speech utterances, daily communication experience appears to shape timing perception, as suggested by the overall bias towards perceiving audiovisual simultaneity when gestures preceded speech, a pattern that mirrors natural timing asynchronies for gestures and the words they semantically correspond to (ter Bekke et al., 2024).

Our findings align with previous evidence of the importance of sensorimotor experience in audiovisual temporal integration. Lee & Noppeney (2011; 2014) demonstrated that musical training selectively influenced the temporal binding of auditory and visual music-related stimuli but not speech-related stimuli, highlighting the impact of experience on audiovisual timing.

However, whereas the previous study examined differences in audiovisual timing sensitivity between participant groups driven by extensive domain-specific expertise (i.e., music versus speech), our study focused on within-subject modulation of audiovisual timing sensitivity within the speech domain itself. In fact, all our participants can be considered speech-gesture experts; nevertheless, we observed enhanced temporal sensitivity specifically for speech-gesture perception patterns shaped by individuals’ own daily communicative experience. This finding suggests that speech-related audiovisual temporal processing benefits not only from long-term, explicit training, but also from implicit, daily experience with one’s own multimodal production patterns.

Interestingly, the TBW was relatively large across all experimental conditions. This result is consistent with previous research, which demonstrates that the TBW is broader for more complex, speech-related stimuli (Stevenson & Wallace, 2013; Van Eijk et al., 2008; Vatakis & Spence, 2006; Wallace & Stevenson, 2014b). Beyond precise timing, temporal flexibility is a crucial skill for effective communication, enabling individuals to adjust to the natural fluctuations in timing and synchrony that occur in real-world interactions (Holler & Levinson, 2019).

The results of this study further refine our understanding of the mechanisms underlying multimodal speech production and those supporting its perception. Although much of the literature has examined audiovisual timing primarily from a perceptual standpoint, the extent to which perceptual and motor timing systems interact, and under what conditions, remains poorly understood. Our study reveals that variability in motor timing predicts perceptual variability in audiovisual speech, but only when participants observe another speaker. This finding suggests a conditional coupling between motor and perceptual timing mechanisms, rather than a strict shared or dissociated system. Given our finding that experience in one domain can influence processing in the other, we hypothesized that the coordination of multimodal speech may rely on mechanisms like those involved in perceiving its temporal structure. This hypothesis is supported by a previous study by Stanczyk et al. (2023) which found a relationship between participants’ performance in a motor timing task and their accuracy in judging temporally asynchronous stimuli.

Interestingly, although motor experience clearly influenced temporal sensitivity in gesture– speech perception, we did not find a direct correlation between variability in gesture-speech production and perceptual binding. This suggests that motor experience alone cannot fully account for how auditory and visual speech signals are integrated. One possible explanation is that shared timing mechanisms support both production and perception, but their influence on audiovisual integration depends on additional factors. Participants appear to possess a well-defined internal model of their own production timing, including its variability. As a result, when observing their own speech, they can reliably predict the onset of the spoken word from gesture initiation, allowing them to maintain a relatively narrow temporal binding window. This interpretation aligns with neuroimaging findings (Bueti et al., 2008; Macar et al., 2006; Merchant et al., 2013; Schubotz et al., 2000) which consistently identify common brain regions involved in both time perception and motor timing, such as the basal ganglia, the cerebellum, the Supplementary Motor Cortex (SMA) and the Premotor Cortex (PMC), while also highlighting the distinct contributions of certain brain areas to separable components of timing.

The findings of this study have significant implications for understanding communication in children who struggle with developing multimodal language skills. For example, autism spectrum disorder (ASD) is often linked to challenges in multimodal integration (Brandwein et al., 2013; Zhou et al., 2018), including difficulties in processing both verbal and non-verbal cues (Smith & Bennetto, 2007). Our results highlight the importance of daily sensorimotor experience in improving perceived speech-gesture timing, with a particular emphasis on the role of the one’s own movements. In individuals with ASD, atypical speech motor patterns (Trayvick et al., 2024) may contribute to difficulties in audiovisual integration when perceiving multimodal communicative acts (Key & Slaboch, 2021), as sensorimotor experience is essential for shaping temporal processing. This raises the possibility that interventions targeting speech motor behavior could indirectly enhance perceptual integration in multimodal communication.

## 4. Limitations

There are several limitations that must be acknowledged. First, our stimuli were restricted to a limited set of sentences and speakers, and the speech was rehearsed rather than spontaneous. This may constrain the generalizability of the findings to more diverse and naturalistic forms of speech and gesture. Second, although our results point to a role of internal models in shaping audiovisual temporal perception, we did not directly measure participants’ explicit awareness of their own production variability or their predictive use of gestural cues.

Future work combining behavioral measures with explicit prediction tasks or computational modelling would help clarify the nature and precision of these internal models. Finally, while our analyses suggest differential contributions of motor variability to perception depending on whether participants observed themselves or others, the neural mechanisms supporting this modulation remain speculative. Neuroimaging or neurophysiological approaches will be necessary to directly test how predictive and timing-related processes are instantiated across production and perception networks.

## 5. Conclusion

This study sheds light on the mechanisms underlying multimodal speech perception, especially in terms of timing. It highlights the importance of sensorimotor experience in shaping how we process and interpret gestures. While sensorimotor theories posit that motor experience can refine perceptual processes through shared internal models (Wolpert, Ghahramani, & Jordan, 1995; Friston, 2010), our findings suggest that this link may generalize across domains and tasks. Future studies could identify shared vs. separate neural correlates for motor and perceptual gesture-speech timing. Such work could help clarify whether, and under what conditions, perception-production coupling emerges in multimodal communication. Understanding how the brain processes and integrates gestures with speech also has broader implications for improving communication in various contexts, such as in educational settings or therapeutic interventions.

